# In plants, decapping prevents RDR6-dependent production of small interfering RNAs from endogenous mRNAs

**DOI:** 10.1101/013912

**Authors:** Angel Emilio Martínez de Alba, Ana Beatriz Moreno, Marc Gabriel, Allison C. Mallory, Aurélie Christ, Rémi Bounon, Sandrine Balzergue, Sebastien Aubourg, Daniel Gautheret, Martin D. Crespi, Hervé Vaucheret, Alexis Maizel

**Affiliations:** Institut Jean-Pierre Bourgin UMR 1318, INRA, SPS Saclay Plant Sciences, Versailles, France.; Institut des Sciences du Végétal, CNRS UPR 2355, SPS Saclay Plant Sciences, Gif-sur-Yvette, France.; Institute for Integrative Biology of the Cell, UMR 9198 - Universite Paris-Sud, Gif-sur-Yvette, France.; Unité de Recherche en Génomique Végétale, UMR INRA 1165-CNRS 8114-UEVE, Evry, France.; Center for Organismal Studies, University of Heidelberg, Germany.; These authors contributed equally to this work.; Current address: Institut Curie, Unité de Génétique et Biologie du Développement, 26 Rue d’Ulm, 75248, Paris Cedex 05, France

**Keywords:** *Arabidopsis*, RNA, P-body, silencing, siRNA, decapping

## Abstract

Cytoplasmic degradation of endogenous RNAs is an integral part of RNA quality control (RQC) and often relies on the removal of the 5′ cap structure and their subsequent 5’ to 3’ degradation in cytoplasmic processing (P-)bodies. In parallel, many eukaryotes degrade exogenous and selected endogenous RNAs through post-transcriptional gene silencing (PTGS). In plants, PTGS depends on small interfering (si)RNAs produced after the conversion of single-stranded RNAs to double-stranded RNAs by the cellular RNA DEPENDENT RNA POLYMERASE 6 (RDR6) in cytoplasmic siRNA-bodies. PTGS and RQC compete for transgene-derived RNAs, but it is unknown whether this competition also occurs for endogenous transcripts. We show that the lethality of decapping mutants is suppressed by impairing RDR6 activity. We establish that upon decapping impairment hundreds of endogenous mRNAs give rise to a new class of rqc-siRNAs, that over-accumulate when RQC processes are impaired, a subset of which depending on RDR6 for their production. We observe that P- and siRNA-bodies often are dynamically juxtaposed, potentially allowing for crosstalk of the two machineries. Our results suggest that the decapping of endogenous RNA limits their entry into the PTGS pathway. We anticipate that the rqc-siRNAs identified in decapping mutants represent a subset of a larger ensemble of endogenous siRNAs.

## Introduction

RNA turnover is an integral part of eukaryotic regulation of gene expression. It regulates RNA abundance and also degrades dysfunctional transcripts that are both a misuse of cellular resources and a source of potentially damaging proteins if translated (for review see (1-5)). Because RNA turnover occurs in all cells, it needs to be tightly controlled to only target specific RNA. Two modifications, the 5’ cap structure and the 3’ poly(A) tail, together with their associated proteins, largely contribute to distinguish a functional mRNA from a dysfunctional transcript, thus protecting mRNAs from exoribonucleases, ensuring mRNAs stability and facilitating translation. Conversely, the absence or removal of the 5’ cap or 3’ poly(A) tail drastically alters the stability of the mRNAs and triggers its degradation. The removal of the cap structure is catalysed by a set of conserved decapping proteins (DCP). In *Arabidopsis thaliana*, DCP1, DCP2 (TDT), DCP5, VARICOSE (VCS) and possibly DEA(D/H)-box RNA helicase 1 (DHH1) constitute the decapping complex (6-9). DCP2 removes the cap, whereas the other proteins likely contribute to mRNA recognition or stimulate decapping. The decapping enzymes are concentrated in cytoplasmic foci called processing (P)-bodies (6, 9-11) and colocalise with the cytoplasmic 5’-to-3’ exoribonuclease XRN4 (12, 13).

Post-transcriptional gene silencing (PTGS) is a conserved eukaryotic mechanism, which destroys RNAs originating from pathogen infection, high levels of transgene expression or select endogenous mRNAs. In plants, PTGS pathways employ SUPPRESSOR OF GENE SILENCING 3 (SGS3), a protein that likely protects RNAs from degradation, and the cellular RNA DEPENDENT RNA POLYMERASE 6 (RDR6) to transform single stranded RNA into double-stranded (ds)RNA, which is subsequently processed into 21-nt small interfering (si)RNAs by the DICER-like enzyme 4(14-16). RDR6 belongs to a family of RDRs that also includes RDR2, which is involved in transcriptional gene silencing (TGS), and RDR1, which has been implicated in virus defense and may antagonize RDR6 activity (17). siRNAs populations guide ARGONAUTE 1 (AGO 1)-dependent mRNA target cleavage through base pairing. RDR6 and SGS3 accumulate in cytoplasmic siRNA-bodies that are distinct from P-bodies (18, 19).

RNA turnover and PTGS are both spatially and functionally linked (20-24). Although it was previously shown that transgene-PTGS is boosted when nonsense-mediated decay, deadenylation, decapping or XRN- or exosome-mediated degradation is impaired (20-24), there is no evidence supporting that endogenous RNAs are processed analogously.

Here, we explore the functional and spatial relationship between decapping and PTGS in *Arabidopsis*. First, we show that mutations in three components of the decapping machinery, DCP1, DCP2 and VCS, provoke an increase of RDR6-dependent transgene PTGS. We also show that a mutation in *RDR6* suppresses the lethality of *dcp2* and *vcs* null alleles. Third, we uncover the existence of a new class of siRNAs (rqc-siRNA) produced by hundreds of endogenous mRNAs upon decapping impairment. rqc-siRNA production from a subset of these mRNAs depends on their RDR6-mediated conversion to dsRNA. Finally, we observe that although appearing as distinct bodies, P- and siRNA-bodies often are spatially associated and display concordant, actin-dependent, movement in the cytoplasm. Together, our data support a model where P-body-localized decapping of endogenous mRNAs deters dysfunctional mRNAs from entering the siRNA-body localized PTGS pathway, and, as such, circumvents the production of rqc-siRNAs, which could direct the sequence-specific degradation of functional cellular mRNAs. These results reveal the importance of a careful balance of RNA turnover and RNA silencing processes for maintaining transcriptome integrity and, consequently, proper plant development.

## Materials and methods

### Plant material and growth conditions

All plants are in the Columbia accession with the exception of the *dcp2^tdt-1^* mutant (25), which has been back-crossed six times to L*er* and which was kindly provided by L.E. Sieburth (8). Lines *L1, Hc1* and *6b4*, and mutants *vcs-6* (SAIL_831_D08), *dcp2-1* (SALK_000519), and *rdr6^sgs2-1^* have been previously described (6, 8, 9, 26). The following mutants were identified in this study: *dcp1-3* (SAIL_377_B10), *vcs-8* (SAIL_1257_H12), and *vcs-9* (SAIL_218_E01). For GUS analyses, plants were grown on Bouturage media (Duchefa) in standard long-day conditions (16 hours light, 8 hours dark at 20-22°C) and transferred to soil after two weeks and grown in controlled growth chambers in standard long-day conditions. *dcp1-3, vcs-8* and *vcs-9* were crossed to *Hc1*, which is in the Col ecotype, while *dcp2^tdt-1^* was crossed to *Hc1/*L*er*, i.e. *Hc1* that had been back-crossed 10 times to the L*er* ecotype. For small RNAs profiling, plants were grown on Bouturage media (Duchefa) in long-day conditions (16 hours light, 8 hours dark) at 15°C and aerial portions were collected 12 dag (days after germination).

### Molecular methods

RNA extraction, RNA gel blot analysis, GUS extraction and activity quantification were described before (20). See supplementary information online for more details.

### Small RNAome profiling and analysis

For each genotype, the aerial portions of 20 12-day-old plants were pooled, and two biological replicates were assayed. RNA samples enriched for small fractions were obtained with mirVana™ miRNA Isolation Kit (#AM1560, Ambion®/Life Technologies Corporation). They were checked for their integrity on RNANano chip, using Agilent 2100 bioanalyzer (Agilent Technologies, Waldbroon, Germany). Small RNA-seq libraries were performed according to NEBNext® Multiplex Small RNA Library Prep Set for Illumina instructions with a different bar code for each sample (#E7300S, New England Biolabs, Inc.). Small RNA-seq libraries are checked for their quality on DNA 1000 chip using Agilent 2100 bioanalyzer (Waldbroon, Germany) before Illumina sequencing (Illumina®, California, U.S.A.). The SmallRNA-seq samples have been sequenced in Single-End (SE) with a read length of 100 bases and a multiplexing rate of 10 SmallRNA-seq libraries on one lane of Hiseq2000 machine to obtained around 15 millions reads/sample. We extracted 18-26nt reads from sequencing files, removed reads mapping the tRNA and rRNA and mapped the remaining reads to the *A. thaliana* genome (TAIR10, perfect matches) using Bowtie2 (V. 2.2.0 –(27)). Read counts for annotated genes were obtained using bedtools (V. 2.17.0 –(28)) in a strand-independent fashion. Intergenic blocks of 18-26 nt RNAs were defined using bedtools as follows: we computed intergenic coverage for each independent library, merged coverage files and extracted all segments with non-zero coverage to produce a unique list of intergenic blocks. Read counts were then obtained for each block and library. Differential expression of annotated genes was estimated using the DESeq package (V 1.16.0 –(29)). For each pairwise comparison, read counts were normalized across all replicates and significant differentially expressed features were determined by a p-adjusted value ≤ 0.05 and a log2 fold change >0 (upregulated) or <0 (downregulated). A locus produce siRNAs from both strands when the ratio of sense to antisense and antisense to sense reads was ≥0.25. Metagene representations were produced by a custom R script. All data were deposited to GEO under the reference GSE65056.

### *Nicotiana benthamiana* and *Arabidopsis thaliana* agro-infiltration

Transient expression in tobacco or in 4-day-old *Arabidopsis thaliana* seedlings were performed as described before (Jouannet:2012hw; 30). See supplementary information online for more details.

### Whole-mount indirect immunofluorescence

Whole-mount immunofluorescence was performed on 5-day-old seedlings according to (31), with some modifications. See supplemental material for detailed protocol.

### Imaging and image analysis

See supplemental material for detailed protocol.

### Plasmid and cloning

See supplementary material for details on the plasmids used and cloning procedures.

## Results

### Viable alleles in the decapping machinery exhibit increased transgene PTGS

An enhancement of RDR6-dependent transgene PTGS by mutations in the P-body-resident enzyme DCP2 was previously described (23). We further examined the functional relationships between decapping and PTGS on transgenes and characterized the consequences of mutations in *DCP1*, *DCP2* and *VCS* on PTGS using the well-characterised *Arabidopsis* reporter lines *L1*, *Hc1* and *6b4*, which all carry a *35S:: GUS* transgene (20, 26, 32). To circumvent the PTGS analysis limitations imposed by the seedling lethality of *decapping* null alleles (6), while simultaneously avoiding the potential for p35S-induced interference (33), we identified partial-loss-of-function 35S-free T-DNA insertion mutants in the promoters, introns or at the 3’ end of the coding sequence. New partial-loss-of-function mutants were identified for *dcp1* (hereafter referred to as *dcp1-3*), and *vcs* (*vcs-8* and *vcs-9*, Supplemental Figure S1). First, we tested the effect of *dcp1-3, vcs-8* and *vcs-9* on line *6b4*, which naturally does not trigger PTGS spontaneously. The absence of GUS mRNA and the presence of GUS siRNAs indicated that *6b4* triggers PTGS in these mutant backgrounds (Supplemental Figure S2A, B). Secondly, the effect of *dcp1-3, dcp2^tdt-1^*(8), *vcs-8* and *vcs-9* on *Hc1*, a line that spontaneously triggers PTGS with 20% efficiency, was tested. Our analysis indicated that *Hc1* PTGS was enhanced in *dcp1-3, dcp2^tdt-1^*, *vcs-8* and *vcs-9* mutants (Supplemental Figure S2C, D) Lastly, we tested the effect of the *vcs-6* seedling lethal allele (6) on *L1*. For this, we took advantage of the fact that although PTGS is triggered in 100% of adult *L1* plants, it is possible to monitor an acceleration of *L1* PTGS when endogenous PTGS suppressors are mutated (34). At five, seven and ten days after germination, silencing was enhanced in *L1/vcs-6* mutant plants compared with *L1* plants (Supplemental Figure S2E). Altogether, these results indicate that decapping prevents or suppresses transgene silencing.

### Mutations in RDR6 rescue the lethality of vcs and dcp mutants

Given the antagonistic effect of decapping (*dcp1*, *dcp2*, *vcs*)(23 and our results) and PTGS (*rdr6*) (15, 16) mutations on transgene PTGS, we postulated that the lethality of decapping mutants could result, at least in part, from the enhanced entry of endogenous mRNAs in the PTGS pathway due to the impairment of decapping activity in P-bodies. Indeed, when the RNA turnover machinery recognizes a dysfunctional transcript only this transcript is eliminated, while homologous functional transcripts remain unaffected. In contrast, PTGS is less specific because it involves the production of siRNAs that guide the degradation of both dysfunctional and functional homologous transcripts. Therefore, when P-body components are mutated, it is possible that dysfunctional endogenous mRNAs enter the PTGS pathway, resulting in the indiscriminate degradation of both dysfunctional and functional homologous mRNAs. On this ground, we anticipated that the lethality of null P-body component mutants could be, at least partly, rescued by null siRNA-body component mutants. To test this hypothesis, we crossed the *vcs-6* null mutant (6), which is seedling lethal, to the *rdr6^sgs2-1^* null mutant (15, 26), which exhibits very mild developmental defects. Whereas *vcs-6* single mutants died at the two-cotyledon stage, *vcs-6 rdr6^sgs2-1^* double mutants developed leaves, stems and flowers (Figure 1A and B). However, they remained smaller than wildtype plants and were sterile, indicating that the *rdr6* mutation does not suppress all of the developmental defects caused by the absence of a functional decapping complex. To extend this finding, we also crossed the *dcp2-1* null mutant Goeres:2007jc, Iwasaki:2007gd, Xu:2006kp} to *rdr6^sgs2-1^*. Whereas *dcp2-1* single mutants died at the two-cotyledons stage like *vcs-6* single mutants*, dcp2-1 rdr6^sgs2-1^* double mutants developed like *vcs-6 rdr6^sgs2-1^* (Figure 1C and D).

**Figure 1:**
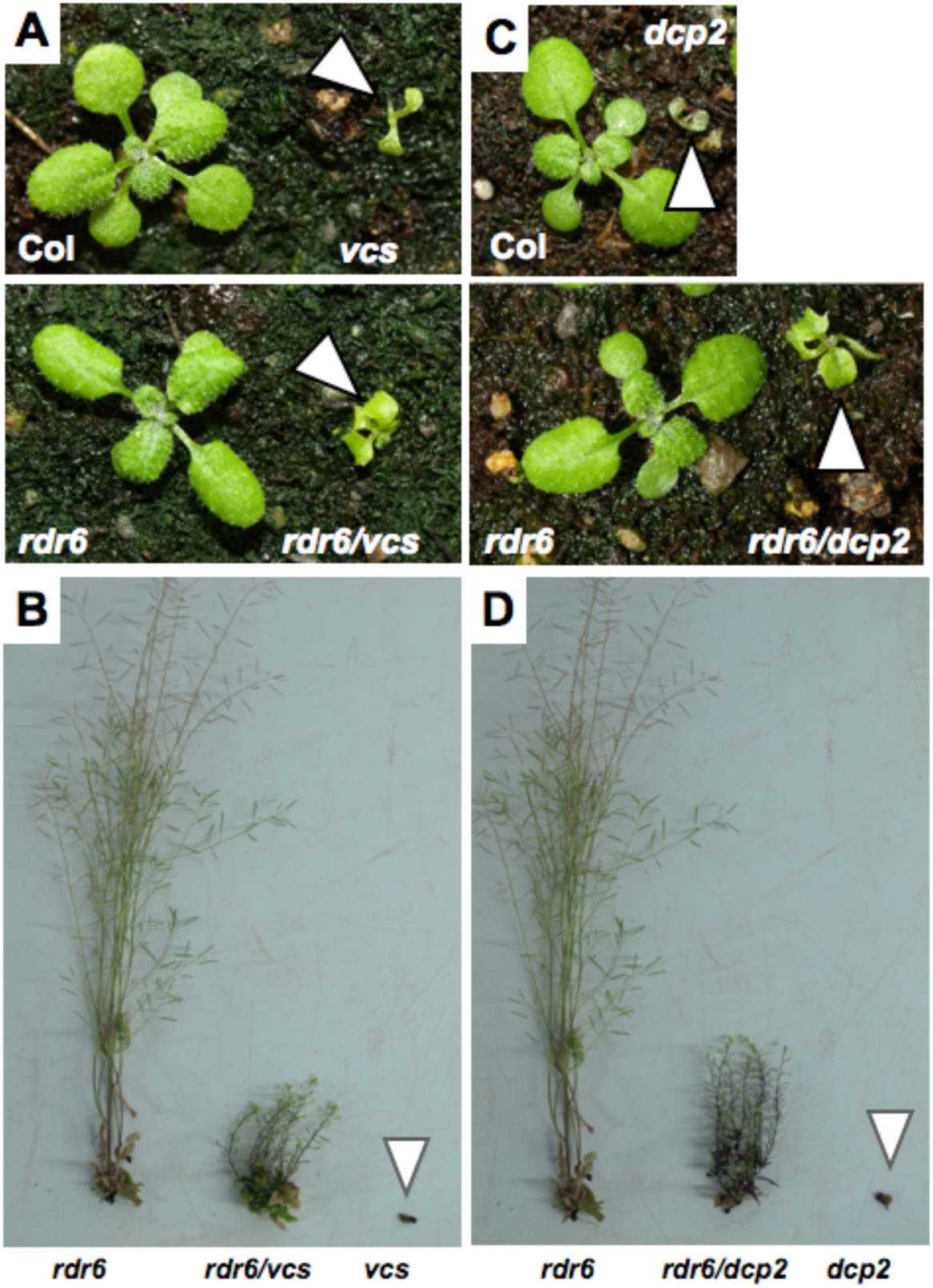
Mutations in *RDR6* allow both the *vcs* and *dcp2* mutants to bypass seedling lethality. Photographs of 15-day-old (A, C) and 65-day-old (B, D) plants of the indicated genotypes. The null mutant alleles *vcs-6* and *dcp2-1* and *rdr6^sgs2-1^* were used.

### Impairing decapping leads to the production of rogue siRNAs from endogenous loci

The *rdr6*-mediated suppression of the lethality caused by the absence of a functional decapping complex, led us to postulate that, in absence of decapping in P-bodies, endogenous mRNAs could enter the PTGS pathway and produce siRNAs. To test this hypothesis, we profiled genome-wide the small (18-26-nt) RNAs present in plants defective in only P-body decapping or plants defective in both decapping and PTGS. We profiled by RNA-seq the small RNAs fractions of two biological replicates of 12-day-old wildtype plants, the null *vcs-6* or *dcp2-1* decapping mutant and the *vcs-6 rdr6^sgs2-1^* or *dcp2-1 rdr6^sgs2-1^* double mutant. For each genotype, the 18-26-nt reads were mapped to the Arabidopsis genome allowing no mismatches to identify transcription units that produce small RNAs (see Supplemental Table S1 and Figure 2). In support of our hypothesis, we identified 1250 transcription units that produced more small RNAs in the *vcs-6* mutant than in wildtype plants (FC>1 and p≤0.05) and 1351 when comparing *dcp2-1* to wildtype (Figure 2A, B and Supplementary Table S3). 816 of these transcription units produced more small RNAs in both *dcp2-1* and *vcs-6* mutants (Figure 2C), indicating that decapping generally prevents the production of siRNAs from hundreds of endogenous loci. We dubbed this new class of small RNAs, rqc-siRNAs because they appear in mutants impaired in RNA quality control. Rqc-siRNAs are predominantly 21 nt long (Figure 2D, E and supplemental Figure S3), and 40% start with a uradine (Figure S4). More than 73% of the loci producing rqc-siRNAs correspond to protein-coding genes (See Supplemental Figure S5). As expected, loci producing rqc-siRNA presented reduced levels of corresponding mRNAs in *vcs-6* mutant but near wildtype levels in the *vcs/rdr6* double mutants (Supplemental Figure S6). For more than 80% of the rqc-siRNAs-producing loci, the rqc-siRNAs are derived from both the sense and antisense strands (Figure 3A, C, E-I). That rqc-siRNAs derive from the two strands of otherwise single stranded mRNAs, suggests that these endogenous mRNAs are transformed to dsRNAs in the *vcs-6* and *dcp2-1* mutants, by an RDR activity. In agreement with this, 20% of the loci producing rqc-siRNAs in *vcs-6* mutant and 15% of those in *dcp2*, produced less rqc-siRNA when *RDR6* is mutated (Figure 2A, B, F and Figure 3B, D, F-I), implicating RDR6 in the production of a subset of these rqc-siRNAs. Our genome wide analysis of small RNA abundance in the *dcp2-1* and *vcs-6* mutants revealed that unlike rqc-siRNAs that become more abundant upon impairment of decapping, the abundance of several miRNAs was reduced in *vcs-6* and *dcp2-1* mutants. 18 and 36 mature miRNAs were respectively reduced in *vcs-6* and *dcp2-1* mutants compared to wildtype (Supplemental Table S4), a result in agreement with prior observations (35).

**Figure 2:**
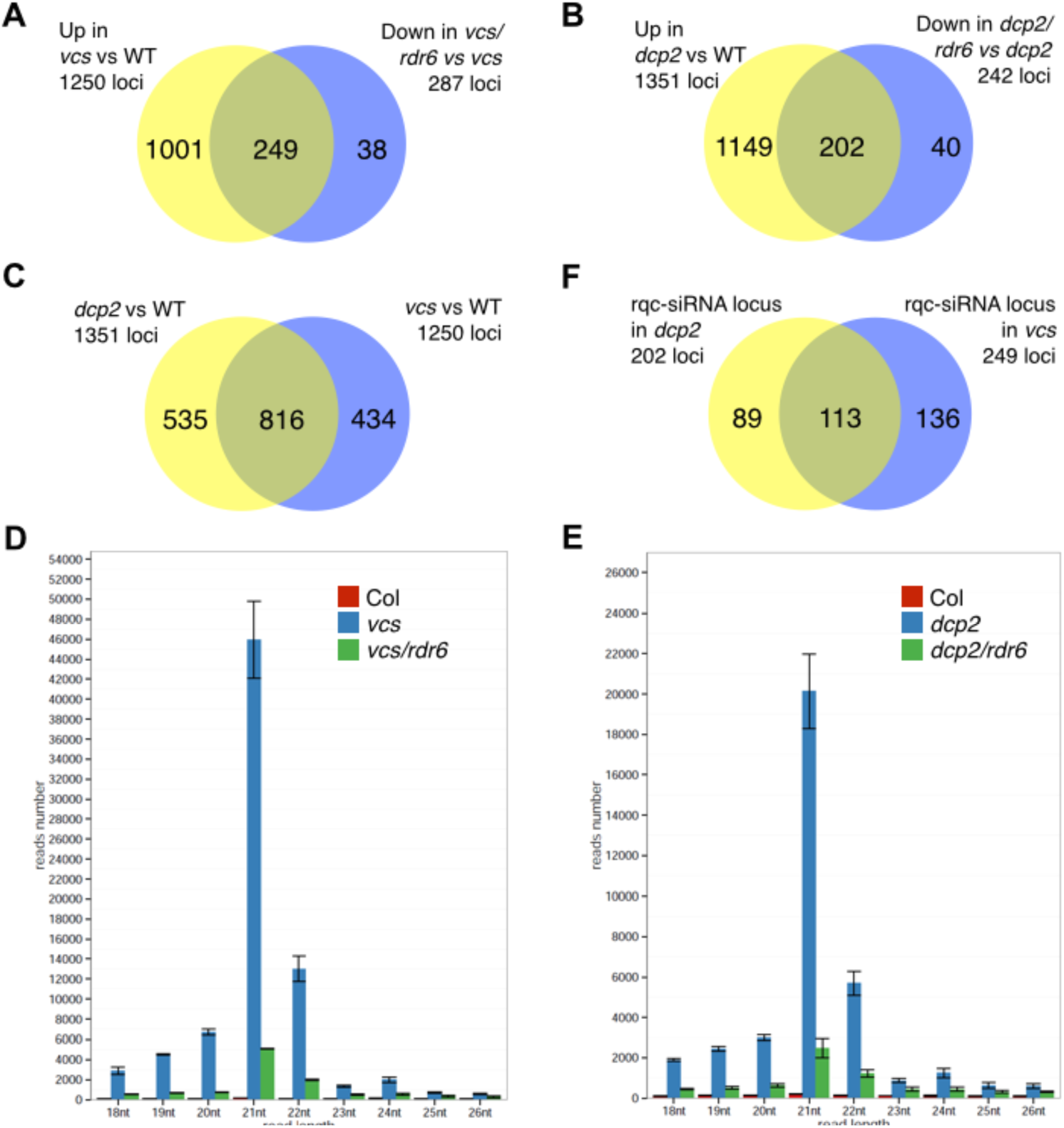
Genome-wide providing of small (18-26-nt) RNAs in plants defective in decapping and PTGS. (A, B, C) Loci with differential accumulation of 18-26-nt small RNAs in wild type, decapping mutants (*vcs-6* or *dcp2-1*) and in double mutants for decapping and PTGS (*vcs/rdr6* or *dcp2/rdr6*). Loci producing more small RNAs (Fold change > 1 and p≤0.05) in (A) or *dcp2-1* (B) mutant compared to wildtype were called “up”, whereas the ones producing less (Fold change < 1 and p<0.05) in the *rdr6/vcs* (A) or *dcp2/rdr6* double mutants were called “down”. The intersection of the two ensemble are the RDR6-dependent rqc-siRNAs. In C, the Venn diagram shows the overlap between rqc-siRNA producing loci in *vcs-6* and *dcp2-1* mutants. (D, E) Size distribution of the RDR6-dependent rqc-siRNAs produced upon impairment of DCP2 and VCS function. Distribution of size (in read number) for the RDR6-dependent rqc-siRNAs produced by the 249 loci upon impairment of VCS (D) and 202 loci upon impairment of DCP2 (E). RDR6-dependent rqc-siRNAs are predominantly 21nt in length. Error bar represents standard error of the mean (sem). In F, is presented the overlap for the subset of rqc-siRNAs dependent on RDR6 for their production.

**Figure 3:**
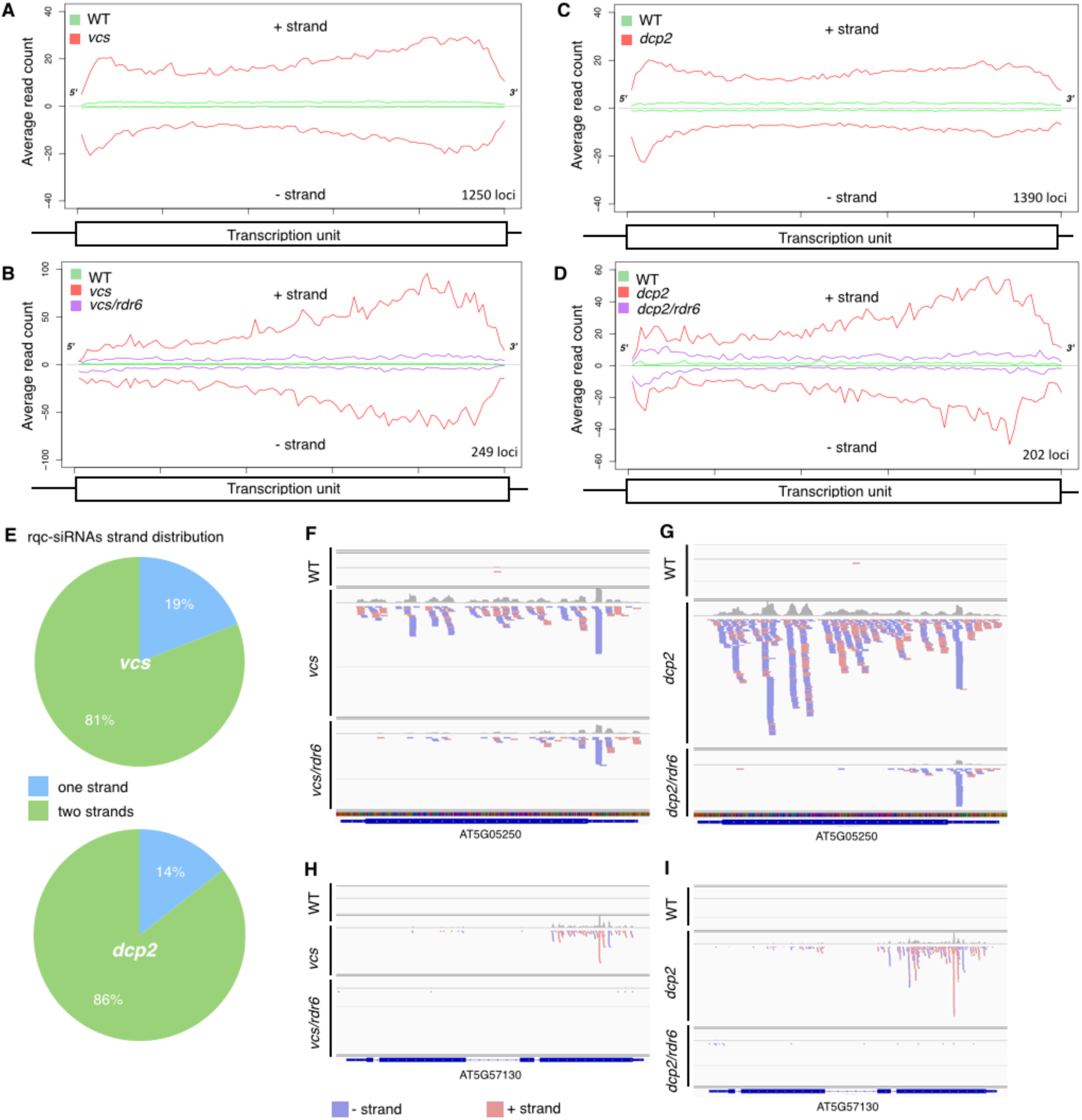
rqc-siRNAs are RDR6-dependent endogenous siRNAs originating from both strands of cellular mRNAs upon impairment of decapping. The average distributions (read count) of 18-26-nt rqc-siRNAs on a canonical transcription unit is represented for the loci that are a source of rqc-siRNAs in the decapping mutants (A, B) and the ones dependent on RDR6 (B, D). (E) Strand distribution of rqc-siRNAs accumulating in *dcp2-1* and *vcs-6* mutants. Most rqc-siRNAs derive from both strands of the producing locus (n=1390 for *dcp2*, n=1250 for *vcs-6*). (F-I) siRNAs distribution on 3 loci producing RDR6-dependent rqc-siRNAs in *vcs-6* and *dcp2-1* mutants. Genome browser views depicting the accumulation of siRNAs in two loci validated by RT-PCR (supplemental Figure S6). (F, H) profiles in wildtype, *vcs-6* and *vcs/rdr6* mutants. (G, I) profiles in wildtype, *dcp2-1* and *dcp2*/*rdr6* mutants.

Altogether, these analyses show that mutations in the decapping complex provoke the production of siRNAs from endogenous mRNAs by the PTGS machinery. This suggests that in the absence of functional decapping enzymes, endogenous RNAs are more prone to becoming substrates for siRNAs production and triggering PTGS. These results also suggest that P-bodies act as a first layer of defense against defective RNAs, preventing or limiting the unintended entry of dysfunctional RNAs into the PTGS pathway, thus circumventing the production of rogue siRNAs that could potentially lead to unintended functional RNAs degradation.

### P-bodies and siRNA-bodies form dynamically linked structures in the cytoplasm

The functional interplay between decapping and PTGS raises the question of where in the cell these two pathways are interacting. To this end, we examined the subcellular localization of decapping and PTGS proteins by expressing translational fusions to fluorescent reporters in *Nicotiana benthamiana* leaves. Coherent with previous studies (6, 13, 18, 19), we observed that DCP1, DCP2, VCS, XRN4, RDR6 and SGS3 fusions accumulated in cytoplasmic foci of two types: RDR6 and SGS3 co-localized in siRNA-bodies, whereas DCP1, DCP2, VCS and XRN4 marked P-bodies (Figure 4A). The accumulation of DCP1, DCP2, XRN4, SGS3 and RDR6 in cytoplasmic foci was confirmed in the root meristem cells of stable *Arabidopsis* transgenic lines expressing GFP-tagged versions of these proteins (Supplemental Figure S8). It was previously reported that P-bodies, marked by DCP1, are distinct from siRNA-bodies, marked by SGS3 (18, 19). We confirmed the existence of these separate foci, although we observed that 50% of the P-bodies and the siRNA-bodies were juxtaposed (Figure 4B, C). The majority of the clusters contained one spot of each class, but association of 3 foci also was observed (Figure 4B, C). We defined as juxtaposed, foci that had no measurable free space between them. The distance from center to center between two adjacent P- and siRNA-bodies was 2.5±1.35μm (average±sd, 89 measurements in 8 independent experiments). The high degree of variation in this measure is linked to the heterogeneity in the size of the foci varying from about 0.2μm to 9μm (mean±sd of 1.87±1.89 μm, n=367). To rule out eventual artefacts due to expression in an heterologous system, we examined the expression of GFP:SGS3 and RFP:DCP1 fusion proteins in *Arabidopsis* cotyledons. We observed that, although distinct, the siRNA- and the P-bodies tended to juxtapose (Figure 4D). This association was further confirmed by whole-mount immunolocalisation of endogenous (untagged) SGS3 in transgenic *Arabidopsis* lines expressing the RFP:DCP1 fusion protein (Figure 4E).

**Figure 4:**
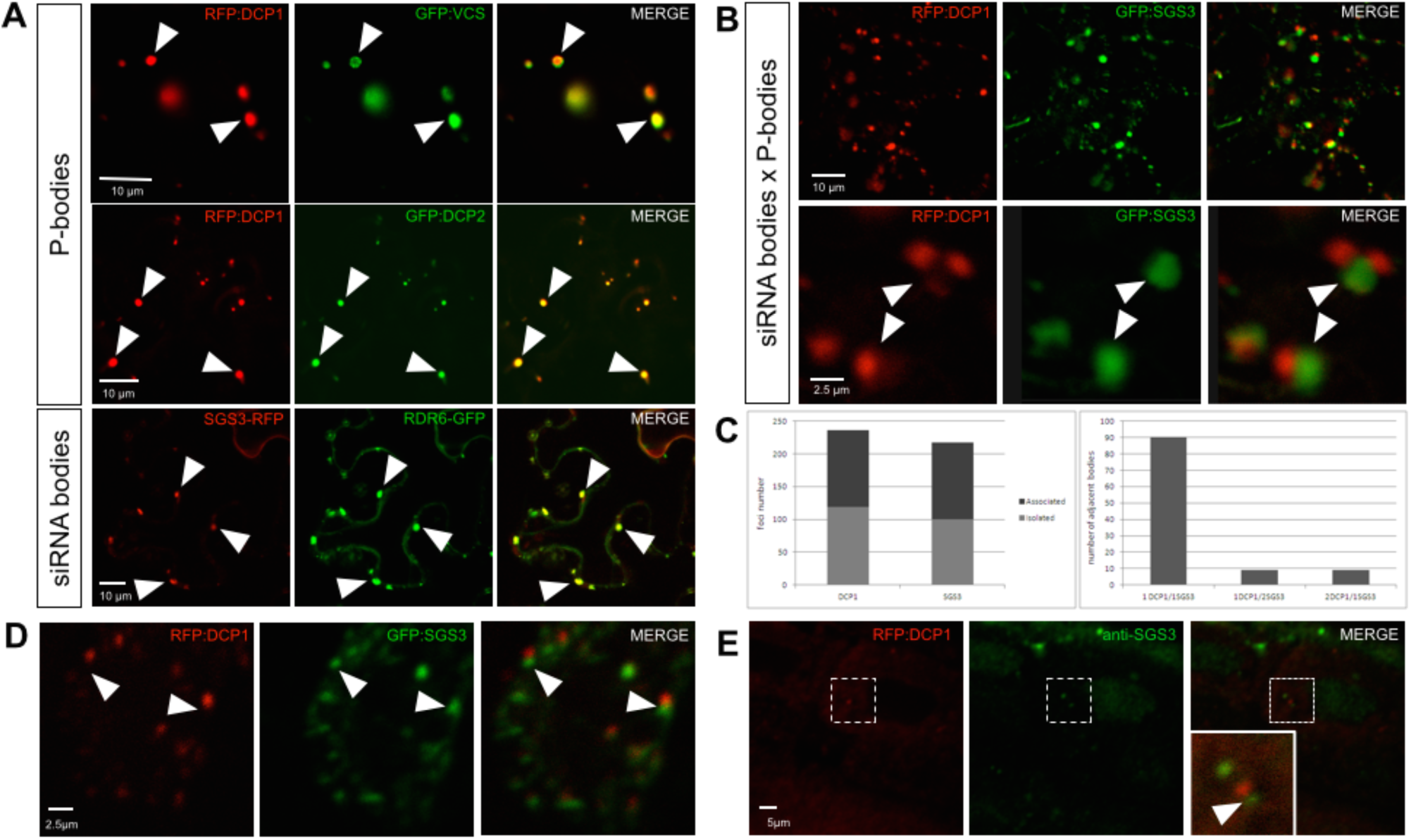
Subcellular localisation of decapping and PTGS proteins. (A) DCP1, DCP2 and VCS accumulate in P-bodies whereas SGS3 and RDR6 accumulate in siRNA-bodies. Confocal sections of *N. benthamiana* leaves co-expressing the indicated fluorescent fusion proteins. The arrowheads indicate the position of a subset of colocalised foci. (B) P-bodies and siRNA-bodies are associated. Confocal sections of a *N. benthamiana* leaf co-expressing RFPDCP1 and SGS3-GFP. The upper row is a low magnification view, and the lower row a close-up on two associated siRNA-/P-bodies. (C) Quantification of P- and siRNA-bodies distribution in *N. benthamiana* leaves expressing DCP1-RFP and SGS3-GFP proteins. Left: quantification of the total number of DCP1 and SGS3 containing bodies found isolated or associated with one another. Right: quantification of the composition of the associated siRNA-/P-bodies (1:1, 1:2 or 2:1). 30 cells in 6 independent experiments were used. (D) Confocal sections of *Arabidopsis thaliana* cotyledons expressing the indicated fluorescent fusion proteins. (E) Whole-mount immunolocalisation of SGS3 in *Arabidopsis thaliana* plants expressing a p35S::RFP:DCP1 construct. In the co-localisation images, the first column represents the RFP or the CFP signal, the middle column represents the GFP or the Alexa488 signal and the third column represents the merged signals for the same plane. In panels B, D and E, the arrowheads indicate the position of the SGS3-marked bodies. Scale bars are shown on the images.

To determine if the observed spatial association between P- and siRNA-bodies reflected transient or more stable association between the two foci, we performed live imaging analysis in *Nicotiana benthamiana* leaves. We observed that the two bodies often remained associated as they moved through the cell (Figure 5A and supplemental Movie S1), indicating an active spatial connection (velocity: 1.12±0.8 μm.s^-1^, mean±sd). We observed several associations and dissociations taking place in short lapses of time between P- and siRNA-bodies (supplemental Movie S2). To characterise this movement further, we monitored P- and siRNA-bodies displacement in the presence of oryzalin, a drug that depolymerizes microtubules, or latrunculinB (latB), a toxin that binds actin monomers and disrupts the actin filaments of the cytoskeleton (36). As previously published, samples were treated for 2h treatment with 10μM of oryzalin or latrunculinB (37, 38). Although mobility of P-bodies has been shown to be dependent on microtubule in animal cells (39), we observed that oryzalin treatment had no effect on the mobility of P- and siRNA-bodies. In contrast, 10μM latB treatment blocked the movement of both bodies (Figure 5B), indicating that in plants the movement of P- and siRNA-bodies depended on actin filaments but not on microtubules.

**Figure 5:**
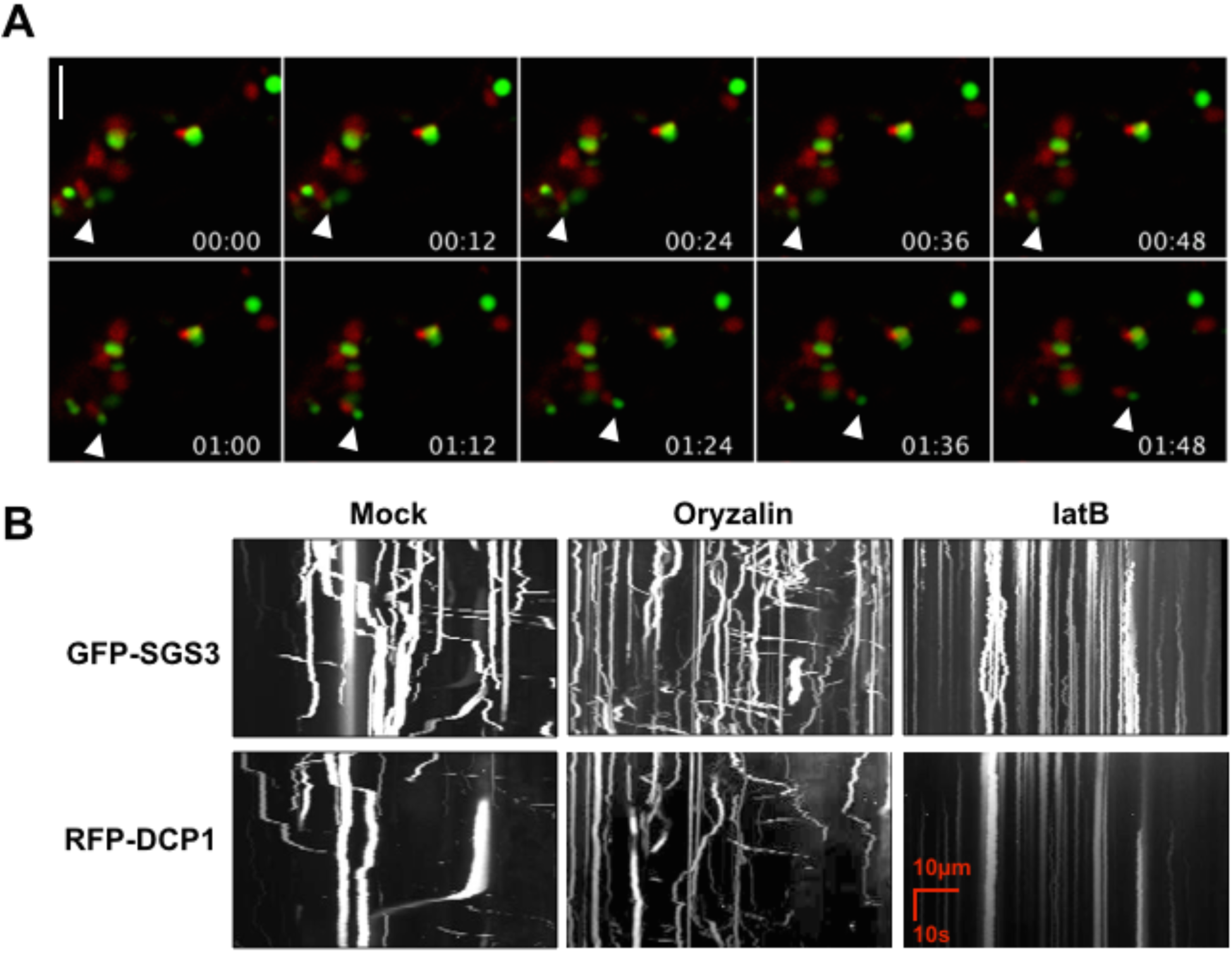
Dynamics of siRNA-bodies and P-bodies. (A) Confocal time lapse recording of a *N. benthamiana* leaf co-expressing RFP:DCP1 and GFP:SGS3. On each frame, the values indicate the time (min:sec) and the arrowheads indicate the displacement of two foci in the cytoplasm over the course of the recording. (B) Kymographs derived from time lapse recordings of *N. benthamiana* leaves co-expressing RFP:DCP1 and GFP:SGS3 and treated either with oryzalin, latrunculin B (latB) or water (Mock). See supplemental material for details. Scale bars 10μm.

## Discussion

In this study, we examined the functional and spatial interplay between P-bodies and siRNA-bodies. We observed that the lethality caused by mutations in the decapping complex components VCS and DCP2 is suppressed by a mutation in the PTGS component RDR6 indicating that these two pathways are linked. Supporting this hypothesis, mutations impairing the decapping step of RNA turnover in P-bodies enhance transgene PTGS and, at the genome-wide level, lead to increased production of rqc-siRNAs from hundreds of cellular mRNAs. Consistent with this functional interplay, we observed P-bodies and siRNA-bodies as distinct foci that dynamically interact in the cytoplasm. We propose that decapping of dysfunctional RNAs in P-bodies prevents their entry into adjacent siRNA-bodies where they could potentially be transformed into deleterious siRNAs.

Our study sheds light on the importance of strict control of the coordination of cytoplasmic exonucleolytic RNA turnover and RNA silencing (PTGS) in plants. When the RNA control machinery recognizes a defective transcript, only the defective transcript is eliminated, while the homologous normal transcripts remain unaffected (*cis*-acting effect). However, under conditions promoting a build-up of aberrant RNAs in the cell, the capacity of the exonucleolytic turnover pathway likely becomes saturated, resulting in the activation of the PTGS pathway. It is reasonable to assume that PTGS is at least as efficient as exonucleolytic RNA turnover in degrading RNAs because PTGS involves the production of siRNAs that guide endonucleolytic RNA cleavage and amplify silencing efficiency, but unlike exonucleolytic RNA turnover, siRNA-guided cleavage does not distinguish a defective transcript from a homologous normal transcript, and thus both types of transcripts would be targeted for degradation (*trans*-acting effect). With PTGS acting in *trans* to degrade all transcripts that are complementary to the siRNA guides, and exonucleolytic RNA turnover degrading individual aberrant/labile transcripts in *cis*, the strict division of RNA substrates between these pathways is likely to be of upmost importance to the integrity of the genome-wide transcriptome (40). Our results suggest that exonucleolytic RNA turnover is the front line defense pathway against defective nucleic acids, while PTGS is induced only when the capacity of the turnover pathways is compromised or saturated. This saturation is consistent with the hypothesis that both a qualitative and quantitative switch in the RNA content triggers PTGS (41, 42). T h i s interplay between PTGS and RNA quality control mechanisms, may extend beyond exonucleolytic RNA turnover. A recent report suggests that non-sense mediated decay (NMD) is a general viral restriction mechanism in plants. Mutation in UPF1 enhances the replication capacity of (+) strand RNA virus, a process counter-balanced by RDR6-mediated PTGS (43). Saturation of NMD by increasing amounts of viral RNAs may constitute a switch for RDR action and siRNA-mediated silencing during infections(43).

Our data provide an additional hypothesis regarding the seedling lethality of decapping mutants(6, 12, 35). It was proposed that lethality of decapping mutants might be caused by the accumulation of a number of different transcripts, indicating that important steps in postembryonic development require decapping-mediated mRNA turnover (6). Alternatively, lethality would be a consequence of the reduced abundance of miRNAs in decapping mutant and/or defects in the miRNA pathway and the subsequent secondary pleiotropic effects on gene expression (35). We propose here that lethality might result from the rqc-siRNA-mediated down regulation of hundreds of cellular mRNA, that become rogue substrates of the siRNA machinery upon impairment of decapping.

We identify rqc-siRNAs as a novel class of siRNAs produced from endogenous loci upon impairment of decapping. In light of the numerous connections between RNA quality control mechanisms and PTGS (20-24), we speculate that the rqc-siRNAs identified here may be a subset of a larger ensemble of endogenous siRNA whose accumulation is prevented by the cellular RNA quality control processes. Impairment of NMD, exosome or deadenylation may reveal the existence of additional rqc-siRNAs producing loci.

The question remains open of what determines an endogenous mRNA to be targeted to the PTGS pathway upon impairment of decapping. We observe significant overlap between the loci becoming source of rqc-siRNAs in the *dcp2* and *vcs-6* mutants. Among these mRNAs, we failed to identify an enrichment for specific GO terms or functional categories when compared to randomly chosen mRNAs (data not shown). Analogously, these transcripts do not present distinctively high levels of expression. The possibility remains open that these mRNAs share common characteristics in their structure or their subcellular localisation.

The rqc-siRNAs produced by endogenous loci when decapping is compromised arise from both strands, a clear signature that they arise from the conversion of the cellular mRNA into a dsRNAs by an RDR activity. That less than 20% of the loci producing rqc-siRNAs in the *vcs-6* or *dcp2-1* mutants required RDR6 suggests that other RDR activities participate in the production of dsRNA in the absence of RDR6. The implication of RDR1 in PTGS (17) suggests that RDR1 could be one such candidate. Recently RDR1 has been shown to be involved in the production of virus-activated siRNAs (vasiRNAs), an abundant class of endogenous 21nt-siRNAs mapped to the exon regions of more than 1,000 cellular genes upon viral infection (44). However, we observe that only 25% of the rqc-siRNAs produced in the *vcs-6* mutant and 12% of the ones produced in the *dcp2-1* mutant are loci also producing RDR1-dependent vasiRNA (Supplemental Figure S7) suggesting that RDR1 may not be an important contributor of the biogenesis of rqc-siRNAs. An alternative scenario could involve overlapping sense-antisense transcription as a source of dsRNAs that would be RDR independent (45).

Several studies have pointed to mRNAs lacking either the 5′ cap or the 3′ poly(A) tail as substrates for RDR6 *in vivo* (22, 46-51). However, *in vitro* studies have shown that purified RDR6 protein is unable to distinguish naked RNAs with a cap structure or a poly(A) tail from naked RNAs that lack one or the other (52). Therefore, it has been postulated that RDR6 is prevented from using intact mRNAs as substrates by the presence of the Cap-binding proteins (CBP), exon junction complexes, and poly(A)-binding proteins (53). This hypothesis is in accordance with our observations that mutants defective in CBP20 or CBP80/ABH1 exhibit enhanced PTGS (34). However, enhanced PTGS in *cbp20* and *cbp80* mutants could also be attributed to a dramatic increase in AGO1 protein level due to the impairment in miR168-mediated *AGO1* repression. Indeed, *cbp20* and *cbp80* mutants have effects on PTGS that are similar to those of *dcl1, hst, hyl1* and *se* mutants, which also exhibit a dramatic increase in AGO1 protein levels due to the impairment of miR168-mediated *AGO1* repression (34). Moreover, if simply the presence of the CBPs prevents RDR6 from using intact mRNAs as substrates, then *dcp1, dcp2* and *vcs* mutations that affect decapping should decrease PTGS. However, we observed the opposite effect on PTGS. Therefore, we propose that aberrant RNAs enter the PTGS pathway and become substrates for RDR6 when the exonucleolytic RNA turnover pathway is compromised, either due to mutations such as *dcp1, dcp2* and *vcs*, or by the saturation of P-bodies with an excess of aberrant RNAs. This proposal is consistent with the previously reported tug-of-war model between RNA quality control and turnover pathways and transgene-PTGS (24), a mechanism also proposed for the triggering of silencing directed against viral RNA (43). It is also supported by our observations that endogenous loci produce siRNAs when decapping is impaired and that P- and siRNA-bodies dynamically interact in the cell, potentially allowing RNAs to be funneled from one body to another.

siRNA-bodies have features in common with stress granules (19) and are often abutting to P-bodies (19, 24). A similar situation has been observed in animal cells. Animal stress granules and P-bodies have related composition and function to their plant counterpart. Live imaging revealed that mammalian P-bodies move toward stress granules, transiently dock and then move on (54). These contacts seem to be maintained when the granules move in the cytoplasm. However, whereas in animal the granules move along microtubules (39), we observe that in plants their movement requires the actin network. In yeast and animal cells, the interaction between different RNA granules has led to the hypothesis that mRNAs and their associated proteins could traffic between the different types of RNA bodies. By extension, one could speculate that the interface between P-and siRNA-bodies controls the transfer of such ribonucleoparticles.

In conclusion, the combination of genetic, cellular and genome-wide small RNAs profiling approaches allow us to propose that exonucleolytic RNA turnover, in general, and decapping, in particular, actively preclude the entry of endogenous dysfunctional RNAs into the PTGS pathway. Hence, these pathways prevent the production of rogue siRNAs that would lead to major deleterious consequences through trans-degradation of their complementary functional cellular mRNAs.

## Supplementary data

Supplementary Data are available at NAR Online: Supplementary Tables 1–4, Supplementary Figures 1–7, Supplementary Movies 1-2.

## Authors contributions

AEMdA performed the genetic analysis of decapping mutants on PTGS with the help of ACM and HV. ABM performed the subcellular localisation studies with the help of AC. AEMdA, RB, SB, MG and DG performed the small RNAs analysis. HV, MDC, ACM and AM designed the experiments. All authors contributed to the analysis of the results. AM, HV and ACM wrote the paper with contribution from all the other authors.

## Acknowledgements

We thank M. Fuchs and L. Bald for their help with RT-PCR, T. Elmayan for helpful discussions, B. Letarnac, H. Ferry and P. Marechal for plant care, N. Bouteiller for help with genotyping, F. Cordelières for the ImageJ plug in “KymoToolBox” and Marc Descrimes for sharing his R script for metagene representation. This work has benefited from the facilities and expertise of the Imagif Cell Biology Unit of the Gif campus which is supported by the Conseil Général de l’Essonne.

## References

1. Shoemaker, C.J. and Green, R. (2012) Translation drives mRNA quality control. Nat Struct Mol Biol, 19, 594–601.

2. Schoenberg, D.R. and Maquat, L.E. (2012) Regulation of cytoplasmic mRNA decay. Nat. Rev. Genet., 13, 246–259.

3. Chen, C.-Y.A. and Shyu, A.-B. (2011) Mechanisms of deadenylation-dependent decay. Wiley Interdiscip Rev RNA, 2, 167–183.

4. Doma, M.K. and Parker, R. (2007) RNA quality control in eukaryotes. Cell, 131, 660–668.

5. Isken, O. and Maquat, L.E. (2007) Quality control of eukaryotic mRNA: safeguarding cells from abnormal mRNA function. Genes Dev., 21, 1833–1856.

6. Xu, J., Yang, J.Y., Niu, Q.W. and Chua, N.H. (2006) Arabidopsis DCP2, DCP1, and VARICOSE form a decapping complex required for postembryonic development. Plant Cell, 18, 3386–3398.

7. Xu, J. and Chua, N.-H. (2009) Arabidopsis decapping 5 is required for mRNA decapping, P-body formation, and translational repression during postembryonic development. Plant Cell, 21, 3270–3279.

8. Goeres, D.C., Van Norman, J.M., Zhang, W., Fauver, N.A., Spencer, M.L. and Sieburth, L.E. (2007) Components of the Arabidopsis mRNA decapping complex are required for early seedling development. Plant Cell, 19, 1549–1564.

9. Iwasaki, S., Takeda, A., Motose, H. and Watanabe, Y. (2007) Characterization of Arabidopsis decapping proteins AtDCP1 and AtDCP2, which are essential for post-embryonic development. FEBS Lett., 581, 2455–2459.

10. Wen, J. and Brogna, S. (2008) Nonsense-mediated mRNA decay. Biochem. Soc. Trans., 36, 514–516.

11. Xu, J. and Chua, N.H. (2011) Processing bodies and plant development. Curr. Opin. Plant Biol., 14, 88–93.

12. Kastenmayer, J.P. and Green, P.J. (2000) Novel features of the XRN-family in Arabidopsis: evidence that AtXRN4, one of several orthologs of nuclear Xrn2p/Rat1p, functions in the cytoplasm. PNAS, 97,13985–13990.

13. Weber, C., Nover, L. and Fauth, M. (2008) Plant stress granules and mRNA processing bodies are distinct from heat stress granules. 56, 517–530.

14. Gasciolli, V., Mallory, A.C., Bartel, D.P. and Vaucheret, H. (2005) Partially redundant functions of Arabidopsis DICER-like enzymes and a role for DCL4 in producing trans-acting siRNAs. Curr Biol, 15, 1494–1500.

15. Mourrain, P., Béclin, C., Elmayan, T., Feuerbach, F., Godon, C., Morel, J.B., Jouette, D., Lacombe, A.M., Nikic, S., Picault, N., et al. (2000) Arabidopsis SGS2 and SGS3 genes are required for posttranscriptional gene silencing and natural virus resistance. Cell, 101, 533–542.

16. Dalmay, T., Hamilton, A., Rudd, S., Angell, S. and Baulcombe, D.C. (2000) An RNA-dependent RNA polymerase gene in Arabidopsis is required for posttranscriptional gene silencing mediated by a transgene but not by a virus. Cell, 101, 543–553.

17. Lam, P., Zhao, L., McFarlane, H.E., Aiga, M., Lam, V., Hooker, T.S. and Kunst, L. (2012) RDR1 and SGS3, Components of RNA-Mediated Gene Silencing, Are Required for the Regulation of Cuticular Wax Biosynthesis in Developing Inflorescence Stems of Arabidopsis. Plant Physiol, 159, 1385–1395.

18. Kumakura, N., Takeda, A., Fujioka, Y., Motose, H., Takano, R. and Watanabe, Y. (2009) SGS3 and RDR6 interact and colocalize in cytoplasmic SGS3/RDR6-bodies. FEBS Lett., 583, 1261–1266.

19. Jouannet, V., Moreno, A.B., Elmayan, T., Vaucheret, H., Crespi, M.D. and Maizel, A. (2012) Cytoplasmic Arabidopsis AGO7 accumulates in membrane-associated siRNA bodies and is required for ta-siRNA biogenesis. 31, 1704–1713.

20. Gy, I., Gasciolli, V., Lauressergues, D., Morel, J.-B., Gombert, J., Proux, F., Proux, C., Vaucheret, H. and Mallory, A.C. (2007) Arabidopsis FIERY1, XRN2, and XRN3 Are Endogenous RNA Silencing Suppressors. Plant Cell, 19, 3451–3461.

21. Gazzani, S., Lawrenson, T., Woodward, C., Headon, D. and Sablowski, R. (2004) A link between mRNA turnover and RNA interference in Arabidopsis. Science, 306, 1046–1048.

22. Gregory, B.D., O’Malley, R.C., Lister, R., Urich, M.A., Tonti-Filippini, J., Chen, H., Millar, A.H. and Ecker, J.R. (2008) A Link between RNA Metabolism and Silencing Affecting Arabidopsis Development. Dev. Cell, 14, 854–866.

23. Thran, M., Link, K. and Sonnewald, U. (2012) The Arabidopsis DCP2 gene is required for proper mRNA turnover and prevents transgene silencing in Arabidopsis. 72, 368–377.

24. Moreno, A.B., Martínez de Alba, A.E., Bardou, F., Crespi, M.D., Vaucheret, H., Maizel, A. and Mallory, A.C. (2013) Cytoplasmic and nuclear quality control and turnover of single-stranded RNA modulate post-transcriptional gene silencing in plants. Nucleic Acids Research, 41, 4699–4708.

25. Deyholos, M.K., Cavaness, G.F., Hall, B., King, E., Punwani, J., Van Norman, J. and Sieburth, L.E. (2003) VARICOSE, a WD-domain protein, is required for leaf blade development. Development, 130, 6577–6588.

26. Elmayan, T., Balzergue, S., Béon, F., Bourdon, V., Daubremet, J., Guénet, Y., Mourrain, P., Palauqui, J.C., Vernhettes, S., Vialle, T., et al. (1998) Arabidopsis mutants impaired in cosuppression. Plant Cell, 10, 1747–1758.

27. Langmead, B. and Salzberg, S.L. (2012) Fast gapped-read alignment with Bowtie 2. Nat. Methods, 9, 357–359.

28. Quinlan, A.R. and Hall, I.M. (2010) BEDTools: a flexible suite of utilities for comparing genomic features. Bioinformatics, 26, 841–842.

29. Anders, S., McCarthy, D.J., Chen, Y., Okoniewski, M., Smyth, G.K., Huber, W. and Robinson, M.D. (2013) Count-based differential expression analysis of RNA sequencing data using R and Bioconductor. Nat Protoc, 8, 1765–1786.

30. Marion, J., Bach, L., Bellec, Y., Meyer, C., Gissot, L. and Faure, J.-D. (2008) Systematic analysis of protein subcellular localization and interaction using high-throughput transient transformation of Arabidopsis seedlings. 56, 169–179.

31. Michael Sauer, T.P. and Eva Benková, J. (2006) Immunocytochemical techniques for whole-mount in situ protein localization in plants. Nat Protoc, 1, 98–103.

32. Béclin, C., Boutet, S., Waterhouse, P. and Vaucheret, H. (2002) A branched pathway for transgene-induced RNA silencing in plants. Curr Biol, 12, 684–688.

33. Daxinger, L., Hunter, B., Sheikh, M., Jauvion, V., Gasciolli, V., Vaucheret, H., Matzke, M. and Furner, I. (2008) Unexpected silencing effects from T-DNA tags in Arabidopsis. Trends Plant Sci., 13, 4–6.

34. Martínez de Alba, A.E., Jauvion, V., Mallory, A.C., Bouteiller, N. and Vaucheret, H. (2011) The miRNA pathway limits AGO1 availability during siRNA-mediated PTGS defense against exogenous RNA. Nucleic Acids Research, 39, 9339–9344.

35. Motomura, K., Le, Q.T., Kumakura, N., Fukaya, T., Takeda, A. and Watanabe, Y. (2012) The role of decapping proteins in the miRNA accumulation in Arabidopsis thaliana. RNA Biol, 9, 644–652.

36. Sampathkumar, A., Lindeboom, J.J., Debolt, S., Gutierrez, R., Ehrhardt, D.W., Ketelaar, T. and Persson, S. (2011) Live Cell Imaging Reveals Structural Associations between the Actin and Microtubule Cytoskeleton in Arabidopsis. Plant Cell, 23, 2302–2313.

37. Samaj, J., Ovecka, M., Hlavacka, A., Lecourieux, F., Meskiene, I., Lichtscheidl, I., Lenart, P., Salaj, J., Volkmann, D., Bögre, L., et al. (2002) Involvement of the mitogen-activated protein kinase SIMK in regulation of root hair tip growth. 21, 3296–3306.

38. Wang, X., Zhang, J., Yuan, M., Ehrhardt, D.W., Wang, Z. and Mao, T. (2012) Arabidopsis MICROTUBULE DESTABILIZING PROTEIN40 Is Involved in Brassinosteroid Regulation of Hypocotyl Elongation. Plant Cell, 24, 4012–4025.

39. Aizer, A., Brody, Y., Ler, L.W., Sonenberg, N., Singer, R.H. and Shav-Tal, Y. (2008) The dynamics of mammalian P body transport, assembly, and disassembly in vivo. Mol. Biol. Cell, 19, 4154–4166.

40. Carroll, B.J. (2011) RNA decay and RNA silencing in plants: competition or collaboration? 10.3389/fpls.2011.00099/abstract.

41. Elmayan, T. (1996) Expression of single copies of a strongly expressed 35S transgene can be silenced post-transcriptionally. 9, 787–797.

42. English, J.J., Mueller, E. and Baulcombe, D.C. (1996) Suppression of Virus Accumulation in Transgenic Plants Exhibiting Silencing of Nuclear Genes. Plant Cell, 8, 179–188.

43. Garcia, D., Garcia, S. and Voinnet, O. (2014) Nonsense-Mediated Decay Serves as a General Viral Restriction Mechanism in Plants. Cell Host and Microbe, 16, 391–402.

44. Cao, M., Du, P., Wang, X., Yu, Y.-Q., Qiu, Y.-H., Li, W., Gal-On, A., Zhou, C., Li, Y. and Ding, S.-W. (2014) Virus infection triggers widespread silencing of host genes by a distinct class of endogenous siRNAs in Arabidopsis. PNAS, 111, 14613–14618.

45. Wang, H., Chung, P.J., Liu, J., Jang, I.-C., Kean, M.J., Xu, J. and Chua, N.-H. (2014) Genomewide identification of long noncoding natural antisense transcripts and their responses to light in Arabidopsis. Genome Res., 24, 444–453.

46. Luo, Z. and Chen, Z. (2007) Improperly Terminated, Unpolyadenylated mRNA of Sense Transgenes Is Targeted by RDR6-Mediated RNA Silencing in Arabidopsis. Plant Cell, 19, 943–958.

47. Parizotto, E.A., Dunoyer, P., Rahm, N., Himber, C. and Voinnet, O. (2004) In vivo investigation of the transcription, processing, endonucleolytic activity, and functional relevance of the spatial distribution of a plant miRNA. Genes Dev., 18, 2237–2242.

48. Montgomery, T.A., Yoo, S.J., Fahlgren, N., Gilbert, S.D., Howell, M.D., Sullivan, C.M., Alexander, A., Nguyen, G., Allen, E., Ahn, J.H., et al. (2008) AGO1-miR173 complex initiates phased siRNA formation in plants. PNAS, 105, 20055–20062.

49. Montgomery, T.A., Howell, M.D., Cuperus, J.T., Li, D., Hansen, J.E., Alexander, A.L., Chapman, E.J., Fahlgren, N., Allen, E. and Carrington, J.C. (2008) Specificity of ARGONAUTE7-miR390 Interaction and Dual Functionality in TAS3 Trans-Acting siRNA Formation. Cell, 133,128–141.

50. Felippes, F.F. and Weigel, D. (2009) Triggering the formation of tasiRNAs in Arabidopsis thaliana: the role of microRNA miR173. EMBO Rep., 10, 264–270.

51. Chen, H.-M., Chen, L.-T., Patel, K., Li, Y.-H., Baulcombe, D.C. and Wu, S.-H. (2010) 22-nucleotide RNAs trigger secondary siRNA biogenesis in plants. PNAS, 107, 15269–15274.

52. Curaba, J. and Chen, X. (2008) Biochemical activities of Arabidopsis RNA-dependent RNA polymerase 6. J. Biol. Chem., 283, 3059–3066.

53. Chen, X. (2008) A silencing safeguard: links between RNA silencing and mRNA processing in Arabidopsis. Dev. Cell, 14, 811–812.

54. Kedersha, N., Stoecklin, G., Ayodele, M., Yacono, P., Lykke-Andersen, J., Fritzler, M.J., Scheuner, D., Kaufman, R.J., Golan, D.E. and Anderson, P. (2005) Stress granules and processing bodies are dynamically linked sites of mRNP remodeling. J. Cell Biol., 169, 871–884.

